# Quantum Entanglement as a Physical Determinant of Genomic Instability in TP53

**DOI:** 10.64898/2026.11.05.764612

**Authors:** Fargam Neinavaie, Diego Santiago-Alarcon, Christina Richards, Joel S Brown, Héctor Mejía-Díaz, Andrew M. Kramer

## Abstract

Mutations in the TP53 tumor suppressor gene are among the most frequent alterations in human cancers, but whether quantum mechanical effects such as DNA entanglement, influence mutation patterns remain unexplored. Here, we applied a coupled quantum harmonic oscillator model to compute von Neumann Entropy (VNE), a proxy for quantum entanglement, across the TP53 sequence. We found that known mutational hot spots exhibited significantly elevated VNE compared to non-hot spot regions, suggesting that local quantum coupling may reflect structural instability or altered repair accessibility. CpG sites, traditionally linked to methylation-driven mutability, did not exhibit elevated VNE, indicating that quantum properties captured by our closed model are distinct from classical biochemical mechanisms. While these computational results raise the possibility that quantum effects could contribute to genomic instability in TP53, we caution that the analysis is limited by the small number of validated hot spots (n=6) and lacks biological validation. our findings are exploratory and should be interpreted carefully. If confirmed in a larger, multi-gene studies with biophysical measurements, a quantum informed framework could open new avenues for predicting mutation hot spots and understanding cancer mutagenesis. We advocate for interdisciplinary efforts bridging quantum physics and oncology to explore these possibilities further.

## Introduction

The TP53 gene, located on chromosome 17p13.1, encodes the p53 protein, a critical transcription factor involved in DNA repair, apoptosis, and cell cycle regulation (Hainaut & Pfeifer, 2016; Levine, 2020). As a key tumor suppressor, TP53 prevents the proliferation of genetically damaged cells, thereby inhibiting tumorigenesis (Lane & Levine, 2010), and its mutations are among the most frequent genetic alterations in human cancers (Hollstein et al., 1991; Kandoth et al., 2013). These mutations frequently cluster within its DNA-binding domain (Cho et al., 1994; Soussi & Béroud, 2001) at specific “hot spots” (Petitjean et al., 2007). While traditional mutational models emphasize biochemical mechanisms like CpG methylation (Hanson & Liebl, 2022) (see Glossary), or oxidative stress (Cadet et al., 2003), they rarely consider how the physical structure of the DNA molecule itself might contribute to mutation susceptibility. The physical structure of the DNA may impinge on the total energy of the system, including both kinetic energy and potential energy which can be quantified even at the quantum level as the Hamiltonian of system.

Quantum mechanical interactions offer a more precise alternative to classical models for analyzing very small objects with low energies, such as DNA molecules (Aarabi et al., 2025). In a classical system, complete knowledge of an object is derived from its position and momentum (Sussman et al., 2015); however, biological environments are highly complex (Ma’ayan, 2017), and cells do not behave as perfect spheres (Tišma et al., 2024). In the quantum approach, the configuration of a particle, such as an electron in a nucleotide, is described by a wavefunction, where the probability density determines the likelihood of finding the object at specific position (Binney & Skinner, 2009). The total energy of the system is represented by Hamiltonian, which accounts for all forces and interactions within the microenvironment (Binney & Skinner, 2009).

Because these quantum properties allow for the modeling of electron cloud fluctuations as waves rather than discrete particles, they have been proposed as potential factors that contribute to DNA mutagenesis (Santiago-Alarcon et al. 2020; Slocombe et al., 2021). One such phenomenon is quantum entanglement, a nonlocal quantum correlation where the states of multiple subsystems, in this study defined as the individual nucleotide bases within a triplet, are interdependent (Ma et al., 2024). In biological systems, correlation between entangled subsystems are revealed through interactions with the surrounding environment rather than through explicit external measurement.

Here, we apply a coupled quantum harmonic oscillator model (Rieper et al., 2010) to evaluate site-specific von Neumann Entropy along the TP53 sequence, with the expectation that higher entropy indicates less energy is required to perturb such DNA regions (Rieper et al., 2010). By determining whether mutational hot spots exhibit distinctive entanglement signatures, we explore a novel biophysical lens for understanding genomic instability. We hypothesize that TP53 mutational hot spots will correlate with regions of significantly elevated VNE, suggesting that local quantum entanglement serves as a biophysical determinant for genomic instability by making these regions more susceptible to physical or chemical transitions.

## Materials and Methods

The human TP53 coding sequence utilized in this study was obtained from the TP53 Database, a comprehensive resource maintained by the National Cancer Institute. The reference sequence corresponds to SwissProt entry P04637 and includes detailed annotations such as exon boundaries and CpG dinucleotide sites. This resource is publicly accessible and can be retrieved via: https://tp53.cancer.gov/tp53seq

Our study employs the coupled quantum harmonic oscillator model proposed by Rieper et al. (2010). In this framework, each nucleotide base is represented as a three-dimensional anisotropic quantum oscillator, corresponding to its electron cloud fluctuation along the x, y, and z axes (Rieper et al., 2010). Adjacent bases interact through dipole-dipole coupling, leading to collective vibrational modes across the DNA segment. We utilize a vibrational Hamiltonian because, at the quantum level, the electron cloud of each base oscillates at specific frequencies; these vibrations are not thermal, but rather the characteristic quantum motions (normal modes) of the electrons themselves (Gutierrez et al., 2009). adjacent bases interact through dipole-dipole interactions between neighboring bases (Rieper et al., 2010).

The vibrational Hamiltonian is constructed by combining local base-specific oscillation frequencies (diagonal elements) with the coupling strength between neighbors (off-diagonal elements). This matrix is then diagonalized to find the system’s normal mode frequencies, which are used to construct the quantum covariance matrix. From this, we extracted a reduced 6×6 covariance matrix for a specific three-base segment (the central nucleotide and its immediate neighbors).

We reproduced the entropy values for all possible central base triplets (A, C, G, T) reported in Table II of Rieper et. al. (2010). The original study focused on a 17-nucleotide strand and did not fully specify the incorporation to the triplet specific effects. Our model expands on the original by adjusting the polarizability tensors of the central base. In this contex, polarizability represents the ease with which a base’s electron cloud is distorted, influencing dipole-dipole interactions with immediate neighbors. These triple adjusted parameters account for local sequence effects, such as π-electron delocalization and stacking forces (Rieper et al., 2010), which are not captured by fixed properties of isolated nucleotides. We emphasize that this approach is a pragmatic model calibration rather than a first-principles derivation; it serves to reproduce entanglement patterns consistent with known biophysical properties, ensuring our computational framework aligns with the benchmarks established by Rieper et al. (2010).

Entanglement was quantified using von Neumann entropy (Vedral, 2002), derived from the symplectic eigenvalues of the reduced quantum covariance matrix of values calculated foreach three-base segment, a central nucleotide and its immediate upstream and downstream neighbors. Within the Rieper et al. (2010) model, entanglement is not treated as a direct causal agent but rather emerges as a property reflecting the electronic correlations between the fluctuating electron clouds of adjacent nucleotides. Similarly, within the present model, entanglement reflects the underlying correlations of the vibrational coupling of the DNA strand. This quantum correlation is quantified by the eigenvalues of the reduced covariance matrix, which represents the probabilities of different quantum states, the specific configuration of energy and position allowed with the triplet’s environment, (Wilde, 2017) with uncertainty measured by their logarithmic contribution (Nielsen et al. 2010). These correlations represent the vibrational coupling of the DNA strand, where the vibration refers to the coordinated displacement of electron densities across the modeled oscillators. These values are determined by the Hamiltonian and covariance matrix construction described in the following section.

To calculate the site-specific von Neumann Entropy (VNE) for a 17-nucleotide DNA segment, the vibrational Hamiltonian was constructed as a real symmetric 3N×3N matrix, with three degrees of freedom per base. Diagonal elements incorporated base-specific trapping frequencies derived from polarizability tensors via 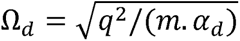, where q is the elementary charge of the electron, m is the effective electron mass associated with the base, and is the base-specific polarizability. Off-diagonal terms encode direction-specific coupling between adjacent bases using the empirical scaling matrix and dipole-dipole interaction formulas from Rieper et al., (2010; see Appendix B, Supplementary Methods for full equations and code). The Hamiltonian was diagonalized to obtain normal mode frequencies (*w_i_*), which were used to calculate the full quantum covariance matrix. A reduced 6×6 covariance matrix was extracted for the central base of interest, and its symplectic eigenvalues were calculated. For readers who wish to see this procedure in action, the full implementation is provided in the R code in the supplementary materials in Appendix B, which can be run to reproduce all intermediate matrices and eigenvalue calculations. The von Neumann Entropy S was then computed using the standard formula for Gaussian states:

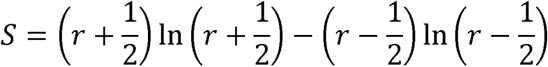

Where r is the mean of the symplectic eigenvalues. Our initial unscaled entropy values systematically underestimated, likely due to incomplete parameter specification in the original theoretical model. We added a uniform scaling factor of ≈5.5. This factor was obtained matching the average unscaled values for adenine-centered triplets in (Rieper et al., 2010). The normalization preserves relative differences between triplets and serves as a practical step in reconstructing entropy values when full parameter details are unavailable.

After calibration, the model was extended to calculate von Neumann entropy values for every position along the TP53 gene. A sliding window of length 3 was moved across the full sequence and for each triplet, the entropy of the central base was calculated using the adjusted quantum harmonic oscillator model. Results for all CpG sites are provided in Appendix C (Table S1).

To assess whether quantum entanglement, as proxied by von Neumann entropy (VNE), varies by genomic context, we compared VNE distributions between CpG sites and non-CpG sites, as well as between known TP53 mutation hot spots and non-hot spots regions. A total of 1171 nucleotide positions across the TP53 gene were analyzed, including 86 CpG sites and 6 mutation hot spots. These hot spots include codons 175 (Krajewska et al., 2003), 245 (Trkova et al., 2003), 248 (Mukhopadhyay & Roth2, 1993), 249 (Peng et al., 1998), 273 (Uddin et al., 2019), and 282 (Baugh et al., 2017), which have been repeatedly identified in tumor sequencing studies. In our statistical analysis, hot spot status was assigned specifically to the individual mutation prone nucleotide within these codons rather than the entire codon triplets. We conducted non-parametric statistical comparison using the Mann-Whitney U test because it does not assume normality and is appropriate for comparing medians between two independent groups, especially when sample sizes differ or when distributions may be skewed (Mann et. al., 1947).

We performed all simulations, entropy calculations, and statistical analyses in R version 4.5.0 (R Core Team, 2025). Key libraries included Matrix version 1.7-3 (Bates, Maechler & Jagan, 2025) for sparse matrix operations. Computations ran on a workstation with an Intel Core i7-9750H processor and 32 GB of RAM running windows 10.

## Results

The overall mean von Neumann Entropy (VNE), used as a quantum mechanical proxy for local entanglement, was 0.0141 (SD=0.00139), with a median value of 0.0140. This narrow standard deviation relative to the mean suggests low variability in entanglement levels at most sites. When stratified by genomic features, CpG sites exhibited a slightly higher mean VNE (0.0147±0.00231) than non-CpG sites (0.0140±0.00128, Figure 1A). The comparison of CpG (n=86) versus non-CpG sites (n=1085) yielded no statistically significant difference in VNE (U=42366, p=0.144), suggesting that CpG status alone does not significantly alter the amount of local entropy.

**Figure 1A.**
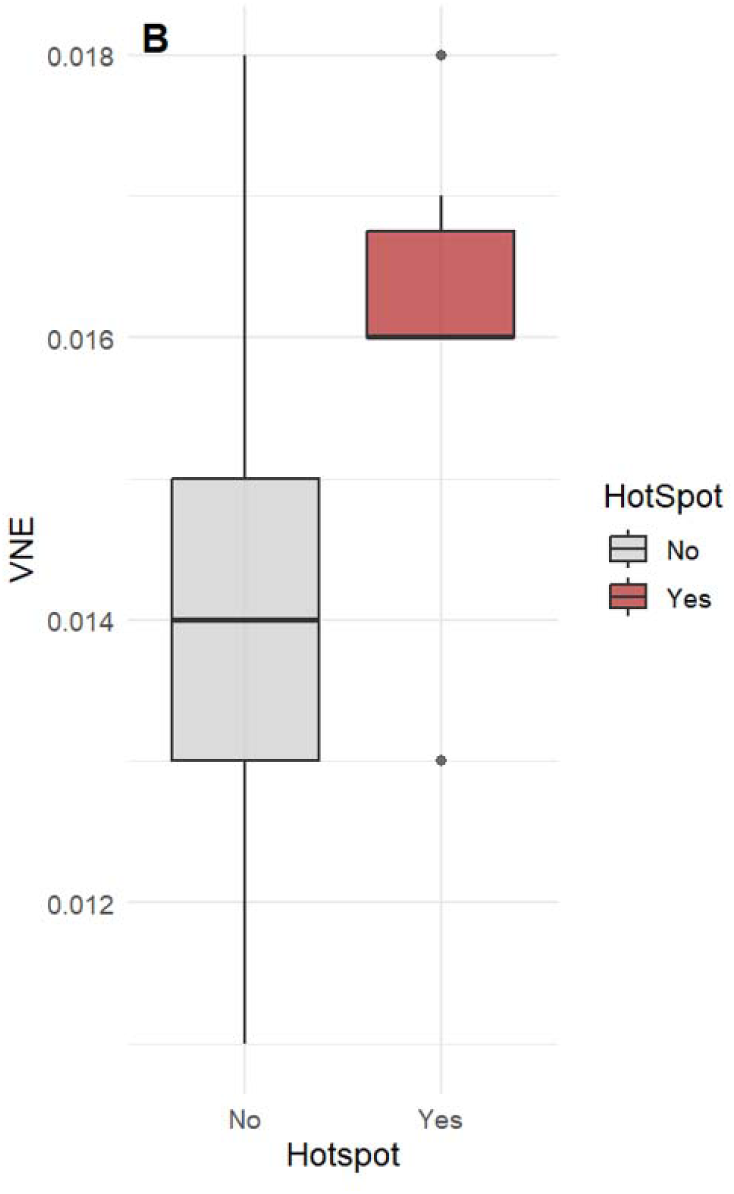
Distribution of von Neumann Entropy by Hot spot Status Boxplot comparing VNE values at mutation hot spots (n=6) a non-hot spot (n=1165) sites. Hot spots exhibit significantly higher median VNE (0.0160 vs. 0.0141; Mann-Whitney U test, p=0.007), suggesting a potential link between quantum entanglement and mutational susceptibility.

Mutation hot spots showed notably elevated VNE values (0.0160±0.00167) relative to non-hot spot regions (0.0141±0.00138). This difference was statistically significant (n=1165) (U=1322, p=0.0069), indicating a measurable association between mutational hot spot status and increased entropy (Fig 1B). Representative VNE values for nucleotides at TP53 mutation hot spot codons are summarized in Table 1.

**Figure 1B.**
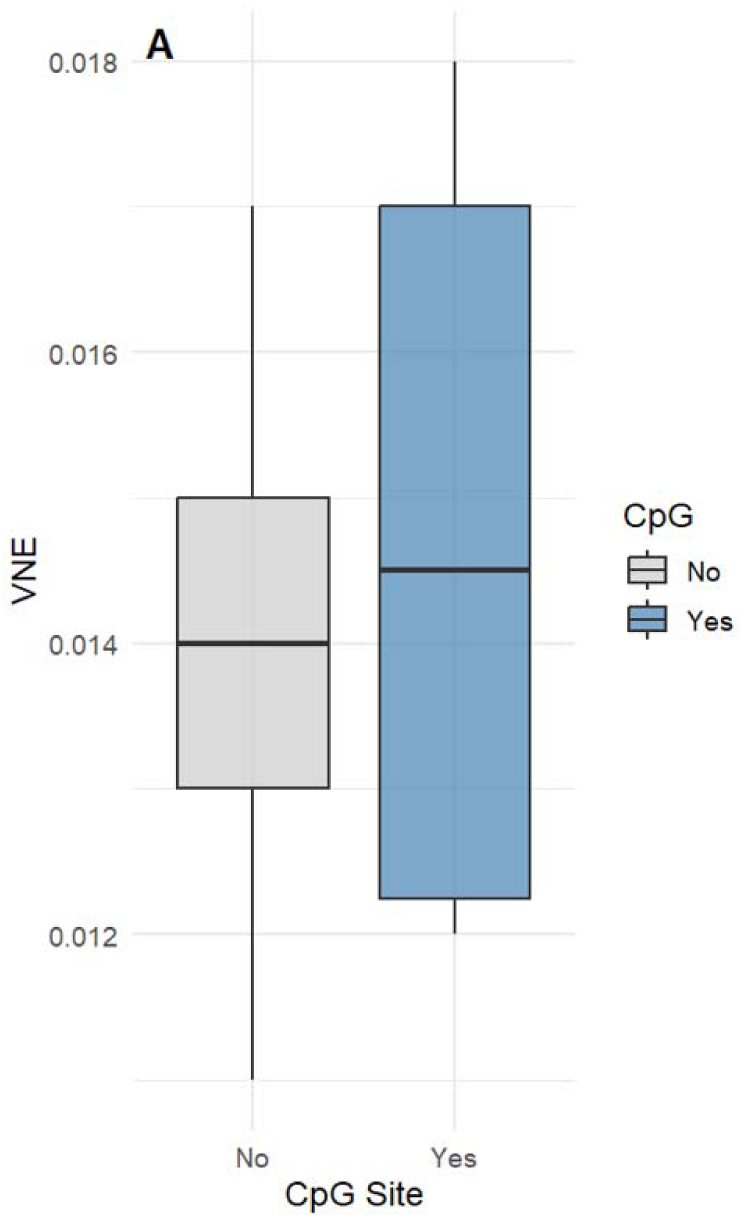
Distribution of von Neumann Entropy by CpG site status Boxplot comparing VNE values at CpG (n=86) and non-CpG (n=1085) sites, CpG sites showed a slightly higher median VNE (0.0147 vs. 0.0140), though the difference was not statistically significant (Mann-Whitney U test, p=0.14).

**Table 1.**
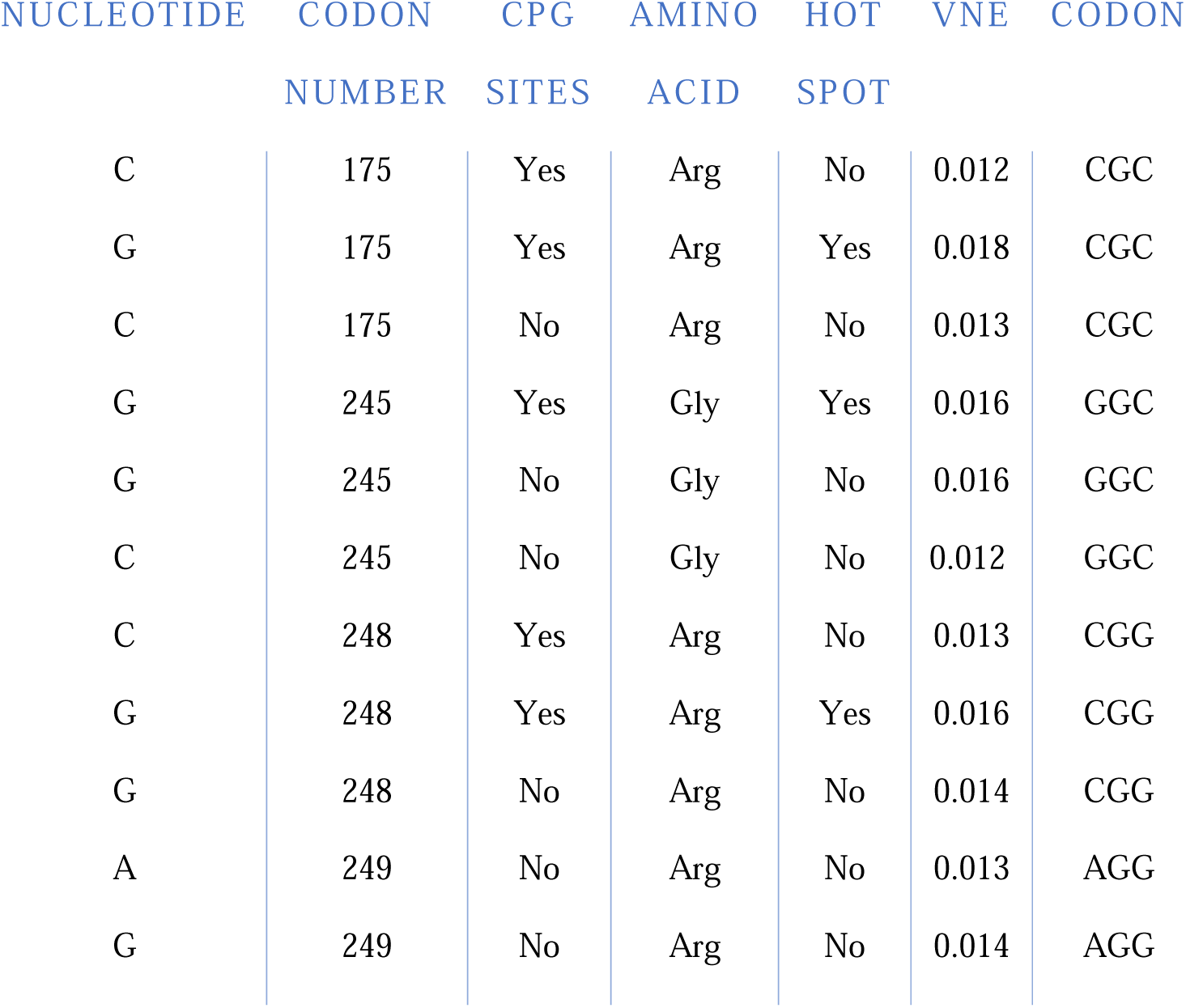
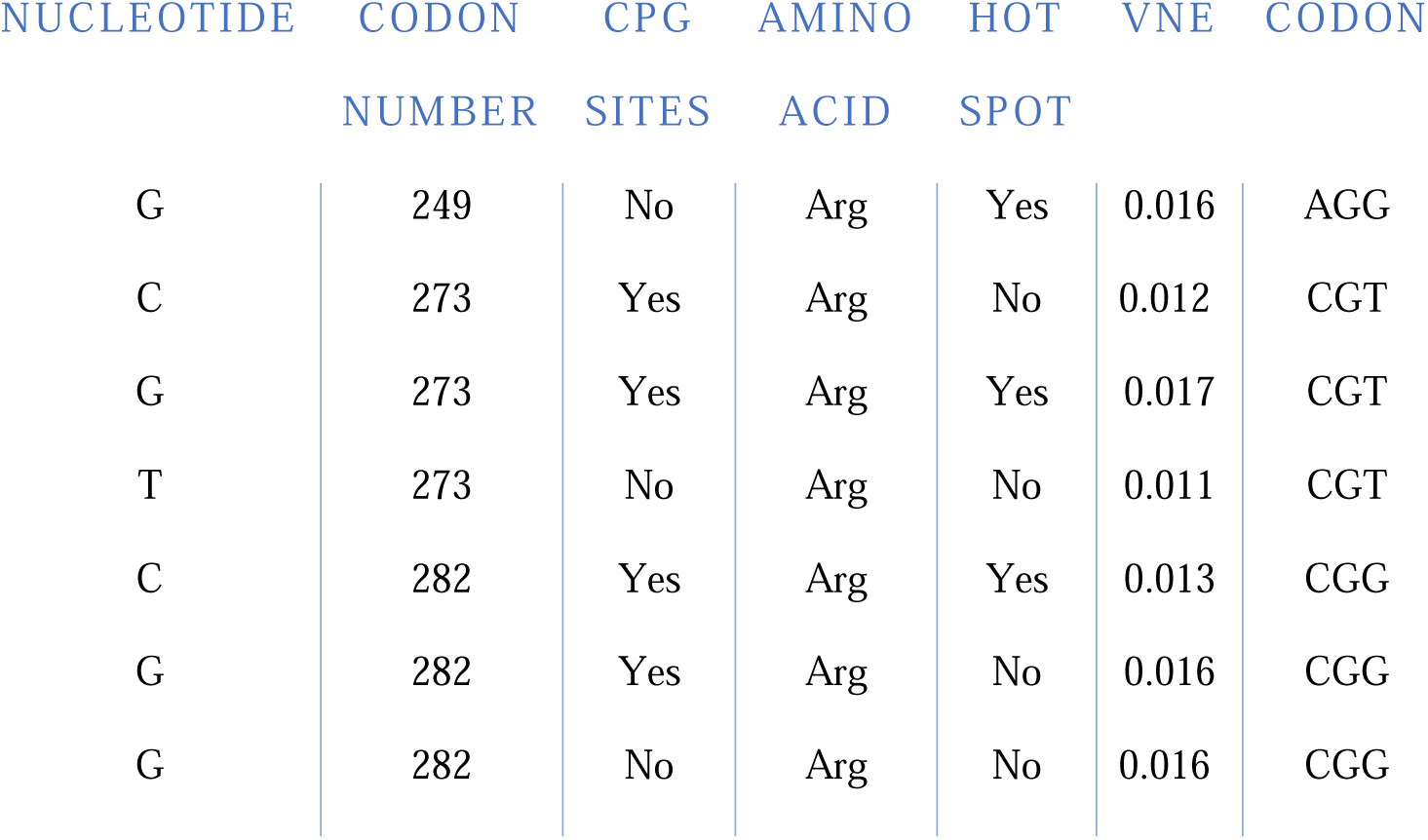
Codon-level characteristics of six TP53 mutation hot spots. Each row represents a nucleotide within the codon of a known TP53 hot spots, showing its identity, CpG status, amino acid, hot spot designation, and corresponding VNE. The hot spot column indicates the specific mutation prone nucleotides (labeled “Yes”) used in the statistical analysis, while neighboring nucleotides within the same codon are labeled “No”.

| NUCLEOTIDE | CODON NUMBER | CPG SITES | AMINO ACID | HOT SPOT | VNE | CODON |
|---|---|---|---|---|---|---|
| C | 175 | Yes | Arg | No | 0.012 | CGC |
| G | 175 | Yes | Arg | Yes | 0.018 | CGC |
| C | 175 | No | Arg | No | 0.013 | CGC |
| G | 245 | Yes | Gly | Yes | 0.016 | GGC |
| G | 245 | No | Gly | No | 0.016 | GGC |
| C | 245 | No | Gly | No | 0.012 | GGC |
| C | 248 | Yes | Arg | No | 0.013 | CGG |
| G | 248 | Yes | Arg | Yes | 0.016 | CGG |
| G | 248 | No | Arg | No | 0.014 | CGG |
| A | 249 | No | Arg | No | 0.013 | AGG |
| G | 249 | No | Arg | No | 0.014 | AGG |
| G | 249 | No | Arg | Yes | 0.016 | AGG |
| C | 273 | Yes | Arg | No | 0.012 | CGT |
| G | 273 | Yes | Arg | Yes | 0.017 | CGT |
| T | 273 | No | Arg | No | 0.011 | CGT |
| C | 282 | Yes | Arg | Yes | 0.013 | CGG |
| G | 282 | Yes | Arg | No | 0.016 | CGG |
| G | 282 | No | Arg | No | 0.016 | CGG |

## Discussion

In this study, we explored the possibility that quantum entanglement, quantified via site-specific VNE derived from a coupled quantum harmonic oscillator model (Rieper et al., 2010), may contribute to the physical underpinnings of mutation prone regions (McFadden & Al-Khalili, 1999; Mejía-Díaz et al. 2025) within the TP53 gene. We found higher entropy at recognized mutational hot spots, indicating lower energy needed for DNA perturbation may be associated with mutations at these sites.

The hot spot at codon 175 demonstrated the highest VNE of all hot spot codons analyzed. Such enhanced coupling may correlate with altered DNA structural stability or with variations in the efficiency of repair enzymes, potentially contributing to the increased likelihood of mutation fixation at these sites. CpG dinucleotides showed a modest, non-significant increase in VNE compared to non-CpG sites. This suggests that the biochemical mechanisms of mutagenesis at CpGs (such as methylation-induced deamination) (Rogozin et al., 2021) may operate independently of the quantum vibrational properties captured by our model, contrasting with the entanglement related susceptibility proposed for non CpG hot spots.

Our model implementation successfully reproduced published entropy values and extended the framework to a real human gene. However, our findings are correlative and do not establish causality. Our adaptation of Rieper et al. (2010) included empirical triplet-based adjustments of nucleotideś electron cloud shifts (i.e., polarizabilities) and a key simplification: we initially employed a 1D harmonic oscillator approximation, whereas the original model operates in 3D. This choice, along with the neglect of nonlinear or long-range interaction, limits the resolution and biological realism of our predictions relative to a full 3D treatment. Future work could integrate molecular dynamics simulations (Yoo et al., 2020), consider chromatin architecture (Wang, 2023), DNA-protein interactions (Ferraz et al., 2021; Winokan et al. 2023), and include environmental factors such as hydration and temperature to enhance biological fidelity (e.g., Mejía-Díaz et al. 2025).

TP53’s mutational landscape is well-characterized (Forbes et al., 2017), with hot spot mutations driving cancer initiation and progression across tumor types (Olivier et al., 2010). Our findings introduce quantum entanglement as an additional factor potentially shaping mutation rates, thus enriching the conceptual framework of genomic instability. If quantum mechanical properties predispose certain loci to mutation, they could serve as predictive markers for hot spot formation and inform models of tumor evolution, including punctuated bursts of mutagenesis (Stephens et al,. 2011) versus gradual accumulation (Alexandrov et al., 2015). Exploring the physical determinant of TP53 mutability could offer insights into vulnerable genomic regions and inform potential strategies for genomic stabilization in clinical settings. Additionally, quantum-derived measures such as von Neumann Entropy (VNE) might serve as hypothetical biomarkers for assessing mutation-related drug resistance or predicting cancer prognosis. An intriguing possibility is that quantum-entangled states, if modifiable through environmental or chemical influences, could one day be targeted to mitigate entanglement-linked instability in key genomic regions.

In a pioneering step at the crossroads of quantum physics and molecular genetics, our study presents computational evidence linking quantum entanglement, quantified through site-specific von Neumann Entropy, with mutational hotspots in the TP53 gene. This work introduces a new quantum biological perspective on genomic instability, underscoring the relationship between DNA’s physical properties and mutation susceptibility. By integrating quantum mechanics with molecular genetics, we open new avenues for understanding cancer etiology and the fundamental forces influencing genome evolution.

## Supporting information

TP53 Gene Sequence

Glossary

Pseudocode: Quantum Entanglement Calculation for DNA Triplets

Table S1. Codon-level von Neumann Entropy values for all CpG sites in TP53.

R code for TP53 analysis

