## Supplementary material for "Quantum Entanglement as a Physical Determinant of Genomic Instability in TP53": Glossary

Appendix A: Glossary

| Term | Definition |
| --- | --- |
| Ab Initio Calculations | Computational methods that use the fundamental laws of quantum mechanics to predict molecular behavior, without relying on experimental data or fitted parameters. |
| Abasic Site | A location in the DNA strand where a nucleotide base is missing, potentially leading to errors during replication or repair. |
| Anisotropic Quantum Oscillator | A quantum model in which vibrations differ depending on direction (x, y, or z axis), representing the electron behavior of DNA bases more realistically. |
| Base Pairing Geometry | The specific 3D arrangement of nucleotide bases (A-T or C-G) when they bond together in the DNA double helix. |
| Base specific Trapping Frequencies | Natural vibration frequencies of each DNA base, influenced by its structure and electron distribution. |
| Cell Cycle Regulation | The cellular control system that manages the timing and order of events in cell division, ensuring genomic stability. |
| Chromatin Architecture | The 3D organization of DNA and its associated proteins within the nucleus, which influences gene expression, DNA repair, and the likelihood of mutations by controlling how accessible different regions of the genome are. |
| Codon | A set of three DNA or RNA nucleotides that codes for a specific amino acid in a protein. |
| Coupled Quantum Harmonic Oscillator | A physics model where each DNA base is treated as a quantum oscillator that interacts with neighboring bases, allowing quantum properties like entanglement to be studied. |
| CpG Dinucleotide | A cytosine followed by guanine in the DNA sequence, often methylated and prone to mutation. |
| Deamination | A chemical reaction in which an amine group is removed from a base (typically cytosine), often resulting in a mutation. |
| Diagonal Elements | Entries in a matrix that represent the self-properties (like vibration) of individual DNA bases, without including interactions. |
| Diagonalization | A mathematical process that simplifies a matrix to find its key characteristics, such as normal modes of vibration. |
| Dipole-Dipole Interaction | A weak physical force between molecules due to the alignment of their electric dipoles; used to model interactions between DNA bases. |
| DNA-binding Domain | The part of a protein (such as p53) that physically attaches to DNA to regulate gene activity. |
| Electron Cloud | The region around a nucleus where electrons are most likely to be found, often modeled as a probability distribution. |
| Entropy (Quantum) | A measure of uncertainty, disorder, or entanglement in a quantum system, used here to represent how “linked” DNA bases are at the quantum level. |
| Genomic Instability | The tendency of a genome to accumulate mutations or structural alterations, often contributing to cancer development. |
| Gaussian State | A type of quantum state where variables like position and momentum follow a bell-curve distribution, common in systems like oscillating molecules. |
| Hamiltonian | A mathematical function that represents the total energy (kinetic + potential) of a system in quantum mechanics. |
| Hot spot (Mutation Hot spot) | A specific DNA location where mutations are observed more frequently than elsewhere. |
| Missense Mutation | A change in a single DNA base that results in a different amino acid being inserted into a protein. |
| Methylation | The addition of a methyl group to a DNA base, which can affect gene expression and mutation risk. |
| Mutagen | A chemical or physical agent that increases the likelihood of mutations in DNA. |
| Mutagenesis | The process by which mutations are introduced into DNA, either spontaneously or through external factors. |
| Mutation | A permanent change in the DNA sequence of an organism, which may or may not affect function. |
| Normal Mode Frequencies | The natural vibration frequencies of a molecule when all parts move in a coordinated way, used here to model base oscillations in DNA. |
| Oxidative Damage | Harm cause to DNA by reactive oxygen molecules. |
| Oxidative Stress | A state in which the balance between harmful reactive oxygen species and protective antioxidants is disrupted, increasing DNA damage risk. |
| π-Electron Delocalization | The spread of electrons across multiple atoms in a molecule, like stacked DNA bases, affecting how the bases interact. |
| Polarizability | A measure of how easily an atom or molecule’s electron cloud can be distorted by an electric filed. |
| Polarizability Tensor | A 3D matrix showing how a base’s electron cloud responds to electric fields in different directions. |
| Quantum Coupling | A situation where quantum systems (like neighboring DNA bases) influence each other’s behavior through shared energy or interactions. |
| Quantum Entanglement | A quantum phenomenon where two or more particles become linked so that the state of one immediately influences the others, even at a distance. |
| Reactive Oxygen Species (ROS) | Unstable molecules containing oxygen that can damage DNA, proteins, and lipids. |
| Reduced Quantum Covariance Matrix | A simplified representation of the quantum fluctuations in just part of a larger system, used here to study a 3-base DNA segment. |
| Replication Error | A mistake made during DNA copying, which can lead to mutations if not corrected. |
| Site-specific von Neumann Entropy | A numerical measure of quantum entanglement calculated for a specific spot in the DNA sequence, based on neighboring base interactions. |
| Stacking Forces | Weak, attractive forces between flat bases in DNA that help hold the double helix together. |
| Stacking Interactions | The way DNA bases align and interact with each other when stacked in the double helix structure. |
| Subsystem | A small part of a larger physical or biological system. For example, one base and its neighbors in DNA. |
| Symplectic Eigenvalues | Special values derived from quantum matrices that are needed to compute entanglement (von Neumann Entropy) |
| Transcription Factor | A protein that binds to DNA and helps turn genes on or off. |
| Transition (Mutation) | A mutation in which one type of base (e.g., C) changes to a chemically similar base (e.g., T) |
| Transversion (Mutation) | A mutation in which a base is replaced with a very different type (e.g., G to T) |
| Triplet-adjusted Polarizability | Modified polarizability values for a DNA base, adjusted based on its neighbors in a three-base segment. |
| Tumor Suppressor | A gene that helps prevent cancer by stopping cells with damaged DNA from dividing. |
| Vibrational Modes | Specific ways in which molecules (like DNA bases) wiggle or vibrate, often used in modeling physical behavior. |
| Von Neumann Entropy | A measure of quantum uncertainty or entanglement, used here to estimate how “connected” or “unstable” a DNA segment may be. |
| Williamson’s Theorem | A mathematical tool used to simplify quantum systems and extract the symplectic eigenvalues needed to calculate entropy. |
