## Supplementary material for "Quantum Entanglement as a Physical Determinant of Genomic Instability in TP53": Pseudocode: Quantum Entanglement Calculation for DNA Triplets

Appendix B: Pseudocode: Quantum Entanglement Calculation for DNA Triplets

1. DEFINE CONSTANTS

- Physical constants (ħ, e, m_e, ε₀, r)

- Base polarizability tensors (α_x, α_y, α_z) for A, C, G, T

- Triplet-adjusted polarizabilities (fit to reproduce Rieper Table II)

- Base-pair coupling scaling matrix (4×4 for A,C,G,T combinations)

2. FUNCTION: build_Hamiltonian(bases)

INPUT: A list of N DNA bases

OUTPUT: A 3N × 3N vibrational Hamiltonian matrix H

FOR each base in sequence:

IF base is center (site 9 of 17):

Use triplet-adjusted polarizability

ELSE:

Use standard base polarizability

Compute oscillator frequency Ω for each spatial direction

Set diagonal H entries to (Ω² × scaling factor)

FOR each pair of adjacent bases:

Compute dipole-dipole coupling strength (based on base identity)

Update off-diagonal terms in H for x-x, y-y, z-z coupling

RETURN H

3. FUNCTION: calculate_triplet_entropy(left, center, right)

INPUT: Identity of left, center, right bases in triplet

OUTPUT: Scaled von Neumann entropy S

Construct 17-base DNA strand with triplet at center (positions 8–10)

Call build_Hamiltonian(bases)

Diagonalize H to obtain ω_i (normal mode frequencies)

Construct position and momentum covariance matrices

Assemble full 6N × 6N covariance matrix

Extract 6×6 reduced covariance matrix for central base

Compute symplectic eigenvalues via Williamson’s theorem

Compute raw von Neumann entropy S using eigenvalues

Apply scaling based on:

- Center base identity

- Neighbor-specific scaling factors

RETURN final entropy value S

4. FUNCTION: generate_central_base_table(center)

INPUT: Fixed central base (A, C, G, or T)

OUTPUT: 4×4 matrix of entropies for all (left, right) combinations

FOR left in {A, C, G, T}:

FOR right in {A, C, G, T}:

S = calculate_triplet_entropy(left, center, right)

Store S in matrix[left, right]

RETURN matrix

5. REPRODUCE RIEPER TABLE

FOR each center base in {A, C, G, T}:

Call generate_central_base_table(center)

Print or store entropy matrix

6. EXTEND TO TP53

INPUT: Full TP53 gene sequence

OUTPUT: Position-specific entropy profile

FOR each position i in 2 to (length - 1):

left = base[i - 1]

center = base[i]

right = base[i + 1]

entropy[i] = calculate_triplet_entropy(left, center, right)

Output entropy profile for visualization and downstream analysis
