## Supplementary material for "Quantum Entanglement as a Physical Determinant of Genomic Instability in TP53": Table S1. Codon-level von Neumann Entropy values for all CpG sites in TP53.

Appendix C: Table S1. Codon-level von Neumann Entropy values for all CpG sites in TP53.

This table includes all codons within the TP53 coding sequence that contain at least one CpG dinucleotide. For each nucleotide in the CpG codons, the base identity, codon position, CpG status, amino acid, hot spot designation, and calculated VNE are reported. While most CpG sites are not mutation hot spots, their entanglement values (VNE) may reflect local quantum properties relevant to mutation susceptibility. This table is provided as a supplement.

| Nucleotide | Codon Number | Amino Acid | Hot Spot | VNE |
| --- | --- | --- | --- | --- |
| C | 4 | Pro | No | 0.013 |
| G | 4 | Pro | No | 0.018 |
| C | 9 | Ser | No | 0.012 |
| G | 10 | Val | No | 0.017 |
| C | 10 | Val | No | 0.013 |
| G | 11 | Glu | No | 0.016 |
| C | 30 | Asn | No | 0.012 |
| G | 31 | Val | No | 0.017 |
| C | 36 | Pro | No | 0.013 |
| G | 36 | Pro | No | 0.017 |
| C | 47 | Pro | No | 0.013 |
| G | 47 | Pro | No | 0.016 |
| C | 48 | Asp | No | 0.012 |
| G | 49 | Asp | No | 0.016 |
| C | 72 | Arg | No | 0.013 |
| G | 72 | Arg | No | 0.018 |
| C | 72 | Arg | No | 0.012 |
| G | 73 | Val | No | 0.017 |
| C | 82 | Pro | No | 0.013 |
| G | 82 | Pro | No | 0.016 |
| C | 83 | Ala | No | 0.012 |
| G | 83 | Ala | No | 0.016 |
| C | 107 | Tyr | No | 0.012 |
| G | 108 | Gly | No | 0.016 |
| C | 110 | Arg | No | 0.013 |
| G | 110 | Arg | No | 0.017 |
| C | 125 | Thr | No | 0.012 |
| G | 125 | Thr | No | 0.017 |
| C | 152 | Pro | No | 0.013 |
| G | 152 | Pro | No | 0.018 |
| C | 153 | Pro | No | 0.013 |
| G | 154 | Gly | No | 0.016 |
| C | 156 | Arg | No | 0.013 |
| G | 156 | Arg | No | 0.018 |
| C | 156 | Arg | No | 0.012 |
| G | 157 | Val | No | 0.017 |
| C | 158 | Arg | No | 0.013 |
| G | 158 | Arg | No | 0.018 |
| C | 158 | Arg | No | 0.012 |
| G | 159 | Ala | No | 0.018 |
| C | 170 | Thr | No | 0.012 |
| G | 170 | Thr | No | 0.016 |
| C | 175 | Arg | No | 0.012 |
| G | 175 | Arg | Yes | 0.018 |
| C | 181 | Arg | No | 0.012 |
| G | 181 | Arg | No | 0.018 |
| C | 185 | Ser | No | 0.012 |
| G | 186 | Gly | No | 0.016 |
| C | 196 | Arg | No | 0.013 |
| G | 196 | Arg | No | 0.016 |
| C | 202 | Arg | No | 0.012 |
| G | 202 | Arg | No | 0.017 |
| C | 213 | Arg | No | 0.013 |
| G | 213 | Arg | No | 0.016 |
| C | 222 | Pro | No | 0.013 |
| G | 222 | Pro | No | 0.018 |
| C | 244 | Gly | No | 0.012 |
| G | 245 | Gly | Yes | 0.016 |
| C | 248 | Arg | No | 0.013 |
| G | 248 | Arg | Yes | 0.016 |
| C | 267 | Arg | No | 0.012 |
| G | 267 | Arg | No | 0.016 |
| C | 273 | Arg | No | 0.012 |
| G | 273 | Arg | Yes | 0.017 |
| C | 282 | Arg | Yes | 0.013 |
| G | 282 | Arg | No | 0.016 |
| C | 283 | Arg | No | 0.012 |
| G | 283 | Arg | No | 0.018 |
| C | 290 | Arg | No | 0.013 |
| G | 290 | Arg | No | 0.018 |
| C | 297 | His | No | 0.012 |
| G | 298 | Glu | No | 0.016 |
| C | 306 | Arg | No | 0.012 |
| G | 306 | Arg | No | 0.016 |
| C | 333 | Arg | No | 0.013 |
| G | 333 | Arg | No | 0.017 |
| C | 335 | Arg | No | 0.012 |
| G | 335 | Arg | No | 0.017 |
| C | 337 | Arg | No | 0.012 |
| G | 337 | Arg | No | 0.018 |
| C | 338 | Phe | No | 0.013 |
| G | 339 | Glu | No | 0.016 |
| C | 342 | Arg | No | 0.013 |
| G | 342 | Arg | No | 0.016 |
| C | 379 | Arg | No | 0.013 |
| G | 379 | Arg | No | 0.018 |
